# Collective cell migration due to guidance-by-followers is robust to multiple stimuli

**DOI:** 10.1101/2023.04.30.538845

**Authors:** Robert Müller, Arthur Boutillon, Diego Jahn, Jörn Starruß, Nicolas B. David, Lutz Brusch

## Abstract

Collective cell migration is an important process during biological development and tissue repair, but may turn malignant during tumor invasion. Mathematical and computational models are essential to unravel the mechanisms of self-organization that underlie the emergence of collective migration from the interactions among individual cells. Recently, guidance-by-followers was identified as one such underlying mechanism of collective cell migration in the zebrafish embryo. This poses the question how guidance stimuli are integrated when multiple cells interact simultaneously. Here, we extend a recent individual-based model by an integration step of the vectorial guidance stimuli and compare model predictions obtained for various variants of the mechanism (arithmetic mean of stimuli, dominance of stimulus with largest transmission interface, dominance of most head-on stimulus). Simulations are carried out and quantified within the modeling and simulation framework Morpheus. Collective cell migration is sfound to be robust and qualitatively identical for all considered variants of stimulus integration. Moreover, this study highlights the role of individual-based modeling approaches for understanding collective phenomena at the population scale that emerge from cell-cell interactions.

## 1 Introduction

Collective cell migration is an important process during biological development, during the morphogenesis of tissues and organs in the embryo, as well as during wound healing and tissue repair in the adult, but collective behavior may also turn malignant during tumor invasion and metastasis [1–4]. Advances in live-cell imaging and precise perturbation experiments allow to study individual cell trajectories and to quantify collective phenomena [5, 6]. Mathematical models and computational analysis are essential to unravel the mechanisms of self-organization that underlie the emergence of collective migration from the interactions among individual cells [7–10]. Integrating experimental and theoretical approaches, different interaction mechanisms between individual migrating cells have been identified, notably chemotaxis during vasculogenesis [11, 12] and during *Dictyostelium* aggregation and slug motion [2], self-generated gradients in the zebrafish posterior lateral line primordium [13, 14], contact inhibition of locomotion during neural crest streaming [15], axis alignment (of polar/ferromagnetic or apolar/nematic type) in local neighborhood [16, 17], chase-and-run between different cell types like neural crest and placodal cells [18], leader cell selection by Notch signaling and guidance *of* follower cells in zebrafish trunk neural crest migration [19] as well as the reverse of the previous mechanism, guidance *by* followers, in polster cell migration toward the animal pole followed by posterior axial mesoderm during zebrafish gastrulation [20]. An important question is how robustly each of these mechanisms can self-organise collective cell migration given variable starting conditions, perturbations and variability of individual cells. Moreover, variability of individual cells may be a prerequisite for leader/follower cell segregation. Potentially, multiple mechanisms need to coexist and synergize to achieve robustness.

The robustness of collective cell migration has been theoretically studied both by individual-based models and continuum approaches [8, 21, 22]. Agent-based or individual-based models (IBM) explicitly describe the stochastic dynamics of individuals and their discrete interactions, in response to their individually differing and dynamic states like current spatial location, movement direction, cell polarity, cell shape or intracellular signaling [8, 23]. A particularly versatile model formalism among IBM is the cellular Potts model (CPM) [12, 24, 25]. Multiple open-source software frameworks allow to develop and simulate CPM in a user-friendly manner [26–28]. Various cell motility modes and all the above mentioned cell-cell interaction mechanisms have been implemented in CPM [10, 29, 30]. To study the robustness of collective cell migration to heterogeneity and fluctuations among cells as well as test alternative model assumptions for a specific interaction mechanism, we will here consider the guidance-by-followers mechanism that coordinates two cell populations in the biological context of early zebrafish development, see Fig. 1.

**Figure 1:**
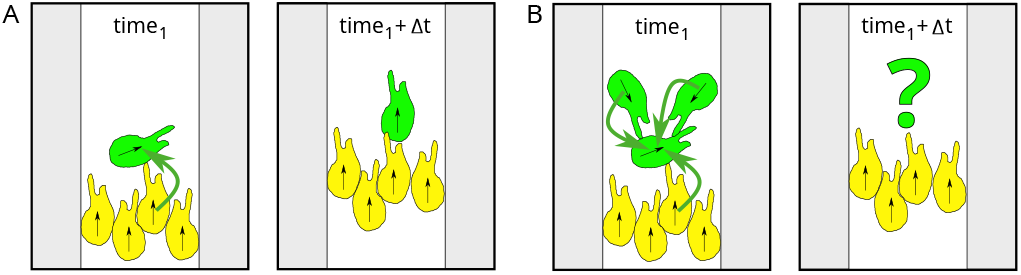
Collective cell migration due to guidance-by-followers as it is underlying polster cell (green) migration followed by posterior axial mesoderm (yellow) during zebrafish gastrulation. (A) Motion direction is transmitted from a moving cell onto another cell (green) when it is hit by the moving cell (here yellow). (B) When multiple cells interact simultaneously, competing input signals need to be processed to determine the direction of motion for each cell. In this study, we propose and compare hypotheses for such signal processing.

During zebrafish gastrulation, the elongating body axis is composed of distinct cell populations. The axis is headed by the mesendodermal polster followed by the posterior axial mesoderm. The constitutive cell types employ different motility modes, polster cells undergo individually random run-and-tumble motility [31] whereas cells of the posterior axial mesoderm advance by convergence and extension [32]. Due to such difference, cell to cell variability and environmental perturbations risk to fragment the body axis. Surprisingly, none of the mutations affecting convergent extension movements and slowing down follower cells (posterior axial mesoderm) leads to axis fragmentation, but rather lead to a slowing of polster cells [32]. Hence, the motility of polster cells is tightly regulated by the left-behind axial cells. At the molecular level, the counter-intuitive propagation of information from follower cells to leading polster cells is mediated by E-cadherin-dependent cell-cell contacts [33] and by E-cadherin/α-catenin mechanotransduction which senses the contact by a moving (follower) cell and regulates front-rear cell polarity and motion orientation [20]. Thereby, the direction of motion can be transmitted from a moving cell onto another cell as it is hit by the moving cell, see Fig. 1A. This fascinating local mechanism is known as ‘guidance-by-followers’ [20]. At the level of the collective, the random cell motility of polster cells gets aligned to form a compact cell cluster in contact with the following posterior axial mesoderm cells. When contact to following posterior axial mesoderm cells gets lost, polster cells no longer receive guidance by posterior axial mesoderm cells and increased randomness of polster cell trajectories then reduces the polster’s net velocity until the following posterior axial mesoderm catches up. Information about the advancement of the posterior axial mesoderm is thereby propagating in the direction of motion faster than the individual cell velocity and polster cells at the front of the axis can therefore respond with net slowing down to perturbations at the rear, as further confirmed by laser ablation and transplantation experiments [20]. In the following, we will call the cells of the posterior axial mesoderm simply axial cells and compare them to polster cells.

Here, our recently developed stochastic and spatially resolved IBM of polster and axial cell interactions, accessible under MorpheusModelID:M0006 (https://identifiers.org/morpheus/M0006) in its previously published implementation [20], is tested against various perturbations. Numbers of cells of the two cell types are varied, cell sizes and therefore contact chances are varied, motility parameters of individual cells are considered heterogeneous and drawn from distributions and motility parameters of individual cells are considered temporally fluctuating. In such simulations, polster cells are exposed to multiple follower contacts at the same time and need to integrate these multiple stimuli, see Fig. 1B. Can competing stimuli mess up collective cell migration? Therefore, we extend this IBM by a processing step of the vectorial guidance stimuli and compare model predictions obtained for three variants of the guidance-by-followers mechanism, (1) arithmetic mean of stimuli, (2) dominance of stimulus with largest transmission interface, (3) dominance of most head-on stimulus. Simulations are carried out and quantified within the modeling and simulation framework Morpheus [28]. Statistics over many repetitions of these stochastic simulations are evaluated. Throughout, collective cell migration from guidance-by-followers is found to be robust and quantitatively identical for all considered variants of stimulus integration, corroborating earlier analysis. Moreover, this study highlights the role of individual-based modeling approaches for understanding collective phenomena at the population scale that emerge from spatially resolved cell-cell interactions.

## 2 Materials and Methods

The model is defined on a two-dimensional hexagonal lattice with periodic boundary conditions and a size of 500 × 1500 grid nodes. There are two static obstacles on either side, leaving a channel of 200 nodes in the middle for the cells to migrate through. These obstacles are included to mimic the observed lateral confinement, which is attributed to paraxial mesoderm. Each grid interval in the simulation represents 1 μm of space, and each time step represents 1 min. The Monte Carlo step duration is set at 0.1 min, allowing for thousands of potential updates per lattice node over the simulated time period. The shape of the cells in the simulations is controlled using a target area of 326 μm^2^, which is an average of 360 cells measured in the experiment [20]. The target circumference is taken from the isoareal circle, favoring circular individual cells if in isolation. Both volume and surface constraints are included in the Hamiltonian with Lagrange multipliers of 1. In addition, the posterior axial mesoderm cells (which are depicted as yellow in the simulations) are given directed motion towards the animal pole (up). The speed of this motion can be adjusted by varying the strength of the ‘DirectedMotion’ term in the Hamiltonian, see the Morpheus documentation [28].

To characterize the cell-autonomous behavior of polster cells (without guidance-by-followers), wild-type polster cells were transplanted to the animal pole of wild-type embryos and migration of isolated cells was tracked [20]. The tracks showed alternating phases of relatively straight migration and phases of slowed and poorly directed movement, indicating that mesendodermal cells exhibit run-and-tumble motion in agreement with earlier observations [31]. To model this behavior, we implemented a run-and-tumble motility with a uniform probability of reorienting the target direction angle, a scaling factor ‘run_duration_adjustment’ for the probabilistic waiting times for reorientation events, which are distributed according to a gamma distribution, and an adjustable Lagrange multiplier that scales the motion speed. The values of the scaling factor and the Lagrange multiplier were obtained via parameter estimation using experimental data of single-cell trajectories [20].

We chose to implement this model as a CPM in the framework Morpheus [28]. Morpheus is an extensible open-source software framework that supports user-friendly declarative modeling. It offers a graphical user interface (GUI) and the domain-specific modeling language MorpheusML to easily compose and extend multicellular models. MorpheusML provides a bio-mathematical language in which symbolic identifiers in mathematical expressions describe the dynamics of and coupling between the various model components. It represents the spatial and mechanical aspects of interacting cells in terms of the CPM formalism and follows the software design rule of separation of model from implementation, enabling model sharing, versioning and archiving in public model repositories. A numerical simulation is then composed by parsing the MorpheusML model definition and automatic scheduling of predefined components in the simulator. Beyond its use in Morpheus, MorpheusML is a model definition standard that enables interoperability among multiple simulators, parameter estimation pipelines and model repositories [34]. Therewith, the FitMultiCell toolbox supports parameter estimation for stochastic Morpheus models (see https://gitlab.com/fitmulticell/fit).

In the model and besides the run-and-tumble motion, mechanical orientation of polster cells can be simulated using the ‘PyMapper’ plugin of Morpheus as described below. For each cell, at fixed time steps of 1 min, the neighbors are detected on 50 membrane points of a ‘MembraneProperty’ data structure and using the ‘NeighborhoodReporter’ in Morpheus. Each cell may possess such a ‘MembraneProperty’ which can be addressed in polar coordinates, is automatically mapped onto the membrane of the cell, moved and deformed along with cell shape changes and allows to execute dynamical models involving multiple ‘MembraneProperty’s as well as coupling to other cell properties. The dynamical model along the membrane may comprise algebraic rules, ordinary differential equations (ODEs) or partial differential equations (PDEs). An example of activator-substrate-depletion pattern formation (two coupled PDEs) on the membrane of a moving cell is accessible from the Morpheus GUI under menu item ‘Examples → Multiscale → CellPolarity.xml’ as well as in the public model repository under MorpheusModelID:M0035 (https://identifiers.org/morpheus/M0035). Instead of PDEs, we’ll here use a rule-based logic to integrate multiple potentially conflicting stimuli.

To implement guidance-by-followers, four scalar ‘MembraneProperty’s store the components of the relative position and velocity direction of each neighbor of a considered cell. The angle between the velocity vector and the direction towards the current cell is then calculated for each neighbor. If this angle is below a threshold called ‘max_angle’ (indicating that the neighbor is moving towards the current cell), the velocity vector of the neighbor is used to update the direction of the considered cell in the ‘DirectedMotion’ plugin, see Fig. 2A.

**Figure 2:**
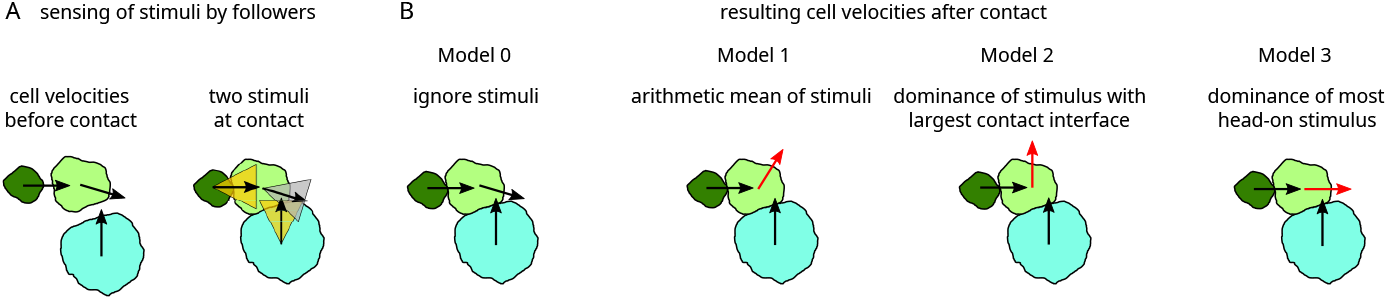
Schematic representation of cell-cell interactions upon contact. (A) Three cells, here of different size, move as indicated by their velocity vectors (left). Upon contact (right), sectors up to the angle *α*_max_ around each velocity vector indicate if guidance is exerted (when the direction to the cell’s center of mass falls within, yellow) or not (grey). (B) Four cell-cell interaction models and their resulting velocity vectors (black unchanged, red changed upon contact): Model 0 is considered as base-line without velocity changes upon contact. Model 1 yields the mean of the impact velocities, weighted by cell-cell contact length. Model 2 yields the velocity vector of the impact with largest cell-cell contact length. Model 3 yields the velocity vector of the impact which is oriented closest to the cell center.

A population of 400 cells is initialized and given 20 min to settle and adjust their shapes and packing. Once this initialization phase is over, two events are triggered by which the cells are assigned an identity based on their position along the anteroposterior (top-down) axis, and the appropriate motility characteristics are applied to them. Unless stated otherwise, we choose to split the population at a position such that one third becomes polster and two thirds become axial cells (see Fig. 3 left). This split establishes a front between polster and axial cells and we will monitor the speed by which this front advances by measuring the average displacement of the first axial cell. The direction in which the front advances will be referred to as frontwards.

**Figure 3:**
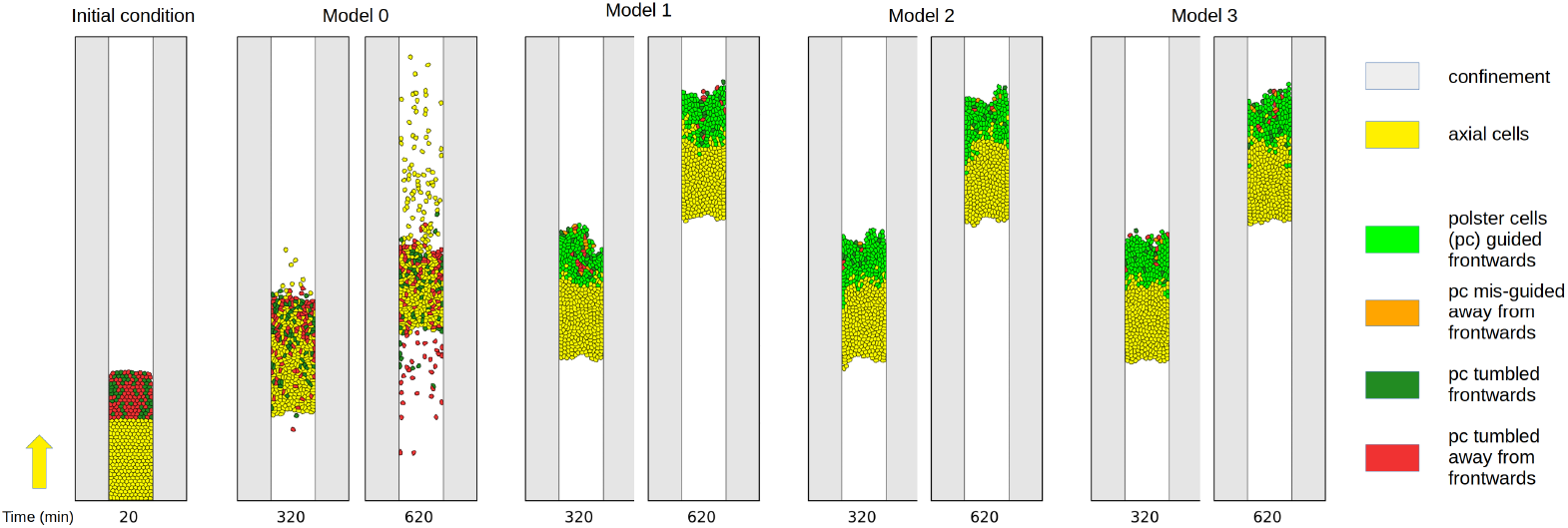
Snapshots of a simulation without the guidance-by-follower mechanism (model 0) and of three different realizations of the guidance-by-follower mechanism (model 1-3) at simulation times 20 min, 320 min and 620 min. Cells are colorcoded based on their identity, source of the velocity vector and alignment. Yellow: Axial cells with motility as indicated by yellow arrow on left. Leading axial cells define the front speed. Light green: Polster cells guided by follower cells such that their orientation points into same quadrant as the axial front velocity. Orange: Polster cells guided by follower cells away from the quadrant of the axial front velocity. Dark green: Polster cells with orientation into same quadrant as the axial front velocity by chance due to run-and-tumble motion. Red: Polster cells with run-and-tumble motion away from the quadrant of the axial front velocity. Visually, all three coupling mechanisms in models 1-3 yield very similar results as opposed to model 0. Simulation time courses are shown in Supplementary Movie 1.

The key model parameters and values as estimated in [20] are given in Tab. 1. If there are multiple neighbor cells moving towards the considered cell, then an additional rule for integrating potentially multiple guidance-by-follower signals is needed, see Fig. 2. To investigate the impact of such signal integration, we here consider and compare four hypothetical signal processing models. In the model implementation, this can be chosen by setting the constant ‘pushing_mode’ to 0-3. The four signal processing models are defined as follows and illustrated in Fig. 2.

**Table 1:**
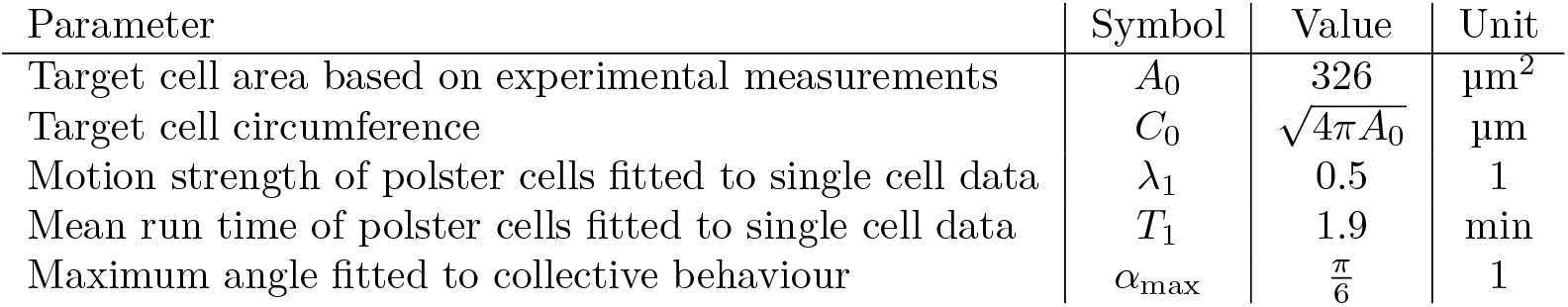
Model parameters with values estimated in [20].

- Model 0 ignores the guidance-by-followers interaction and serves as a quantifiable base line where cells will collide and squeeze past each other due to steric interactions.
- Model 1 sets the cell velocity to the arithmetic mean of neighbors’ velocity vectors which fall within a max_angle-sector, with the length of each cell-cell contact serving as the weighting factor.
- Model 2 sets the cell velocity to that of the neighbor which has the largest contact length given that its velocity vector falls within the ‘max_angle’-sector around the contact vector, ignoring other candidates. If the largest contact does not fulfill the ‘max_angle’-sector criterion because it strikes more tangential, then the next smaller contact is evaluated.
- Model 3 sets the cell velocity to that of the neighbor with the best aligned velocity vector, i.e. smallest angle between velocity and contact vectors, ignoring other candidates.

These hypotheses serve as abstractions of the microscopic biochemical and biophysical mechanism. They yield distinct behaviours for the test configuration in Fig. 2. The Morpheus model generated during this study was deposited in the public model repository under MorpheusModelID:M0008 (https://identifiers.org/morpheus/M0008).

## 3 Results

### 3.1 Robustness to multiple conflicting stimuli

First, we demonstrate that multiple cells can collide simultaneously and that the direction signal is robustly transmitted from axial cells to polster cells and among polster cells for models 1-3, see Fig. 3. Qualitatively, snapshots of the simulations already suggest that all three coupling models 1-3 yield very similar results as opposed to the base line model 0 that just possesses steric interactions. To analyze this in detail, we have varied the motion strength of axial cells between zero and nominal value (Tab. 1) in 5 steps and run 100 repetitions per parameter combination. For each run, we have quantified the angular coherence of polster cell orientation, defined as fraction of polster cells moving in the frontward quadrant (± 45° around the vertical front velocity of axial cells), and the speed of the axial cell front, defined by the anterior-most axial cell, as shown in Fig. 4. The dependency of these two observables can be compared to our previously published experimental data from [20]. For the experimental data points as well as for the simulation data, a linear regression was calculated, see Tab. 2, and plotted along with the prediction error (68 % interval) in Fig. 4. The differences between the coupling models 1-3 were found to be negligible compared to the base-line model 0. Within error margins, all three coupling models can explain the experimental data equally well (Fig. 4). Individual data points of all simulation runs are shown in Supplemental Fig.S1.

**Figure 4:**
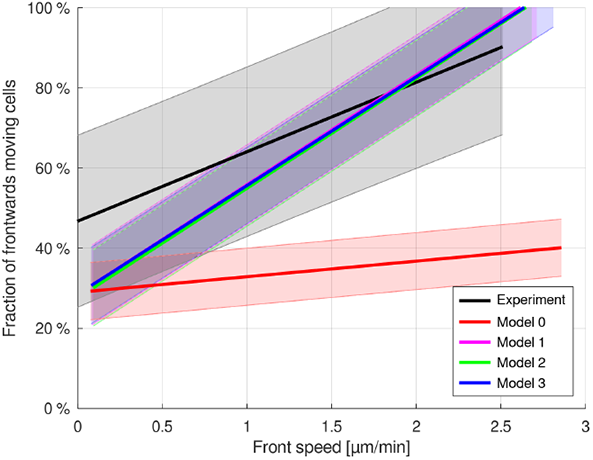
Quantification of angular coherence between guided and follower cells. All three coupling mechanisms yield very similar results and, considering the error margins, are able to explain the experimental data as opposed to Model 0 without guidance-by-followers. Lines show linear regression of the data and shaded bands are the 68 % intervals of the prediction error. Individual data points of all simulation runs are shown in Supplemental Fig.S1.

**Table 2:** Summary of simulation results compared to experimental data by [20]. Each simulation study (figure) comprises 100 runs for each of five different parameter values of axial cell motion strength. Front speed and fraction of frontwards moving cells were quantified for each run and the linear regression coefficients reported in this table were calculated from the 500 data points per model and simulation study.

| Study, Figure | Model | Slope | Offset | Std |
| --- | --- | --- | --- | --- |
| Experimental data, Fig. 4 |  | 17.34 | 46.75 | 13.69 |
| Original parameter values, Fig. 4 | 0 | 3.89 | 28.99 | 4.70 |
|  | 1 | 27.35 | 28.53 | 6.56 |
|  | 2 | 27.39 | 27.58 | 6.27 |
|  | 3 | 27.14 | 28.52 | 6.30 |
| Polster-axial cell ratio 1:5, Fig. 5a | 0 | 5.53 | 27.49 | 6.37 |
|  | 1 | 24.73 | 30.52 | 7.51 |
|  | 2 | 23.87 | 31.00 | 8.09 |
|  | 3 | 24.58 | 29.33 | 7.86 |
| Polster-axial cell ratio 5:1, Fig. 5b | 0 | 1.44 | 27.37 | 2.95 |
|  | 1 | 19.84 | 27.01 | 7.89 |
|  | 2 | 13.75 | 28.16 | 7.04 |
|  | 3 | 14.03 | 29.87 | 6.97 |
| Random cell volume, Fig. 6 | 0 | 3.47 | 29.70 | 4.85 |
|  | 1 | 27.16 | 29.04 | 6.42 |
|  | 2 | 27.04 | 28.32 | 6.58 |
|  | 3 | 27.20 | 28.51 | 6.41 |
| Random motion strength, Fig. 7 | 0 | 3.85 | 29.20 | 4.87 |
|  | 1 | 26.03 | 30.25 | 6.86 |
|  | 2 | 26.42 | 28.69 | 6.96 |
|  | 3 | 26.32 | 29.27 | 6.81 |
| Time-varying motion strength, Fig. 8 | 0 | 4.08 | 29.38 | 4.67 |
|  | 1 | 27.79 | 28.09 | 6.57 |
|  | 2 | 26.85 | 28.02 | 5.89 |
|  | 3 | 27.44 | 27.18 | 6.41 |

### 3.2 Robustness to cell type ratio

In order to test whether the coupling mechanisms are sensitive to the cell type ratio, simulations with ratios between polster and axial cells of 1:5 and 5:1 were conducted (Fig. 5, Tab. 2). In the case 1:5 with abundant axial cells, the number of additional axial cells in the back of the front has no additional effect and front movement is unchanged (Fig. 5a) compared to the above cases with 1:2 ratio because only the axial cells near the front can exert their orienting effect on polster cells. In the case 5:1 with few axial cells, simulations show increasing disorder of the front between axial and polster cells and disordered front movement (Fig. 5b). Movies of the simulations reveal that the abundant polster cells invade the group of axial cell and cause the disorder, thus disturbing the collective effect of guidance-by-followers.

**Figure 5:**
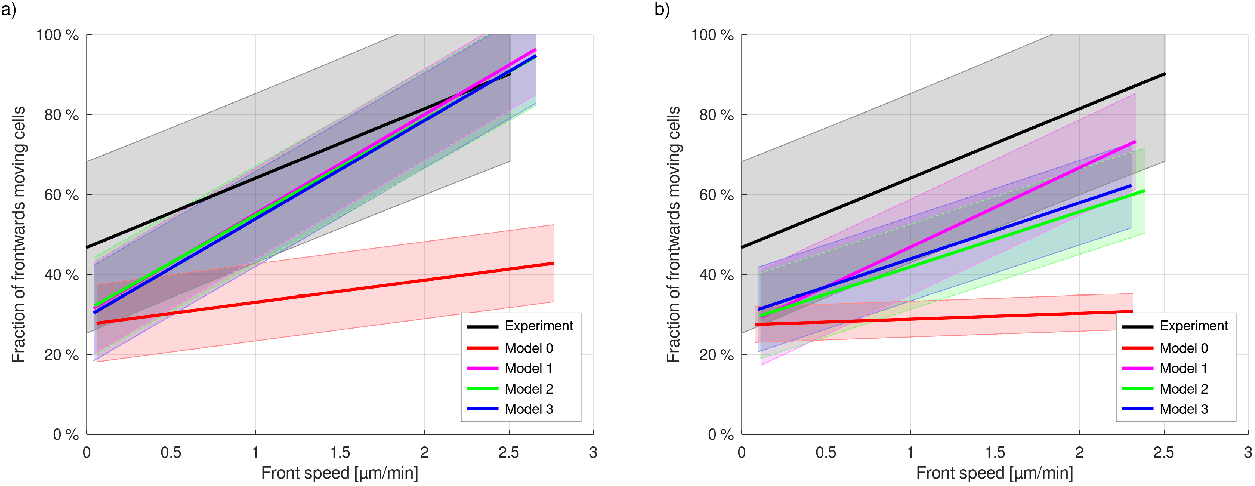
Polster cell to axial cell ratio 1:5 (a) and 5:1 (b). A high polster cell fraction leads to less ordered front cell movement.

### 3.3 Robustness to cell size variability

If cell volumes are not equal, but distributed uniformly across a range 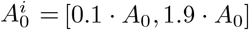 symmetrically around the originally identical cell volume *A*_0_, then practically no deviation from the original behaviour is observed (cf. Figs 6 and 4). This shows that the collective cell migration behaviour is very robust against variability and changes in cell volume.

**Figure 6:**
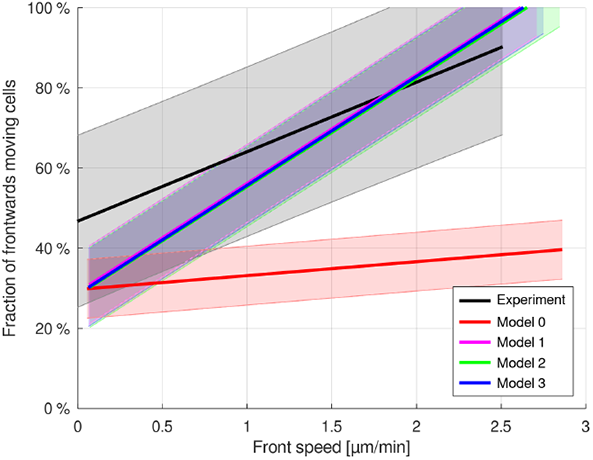
Uniformly distributed cell volumes 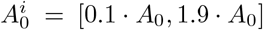 don’t disturb the guidance-by-follower mechanism.

### 3.4 Robustness to random motility parameters

For a uniform distribution of the polster cells’ individual motility strength parameter 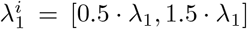 symmetrically around the former strength parameter *λ*_1_, no significant difference was observed (Fig. 7), hence rendering the mechanism very robust against variation in the polster cells’ motility.

**Figure 7:**
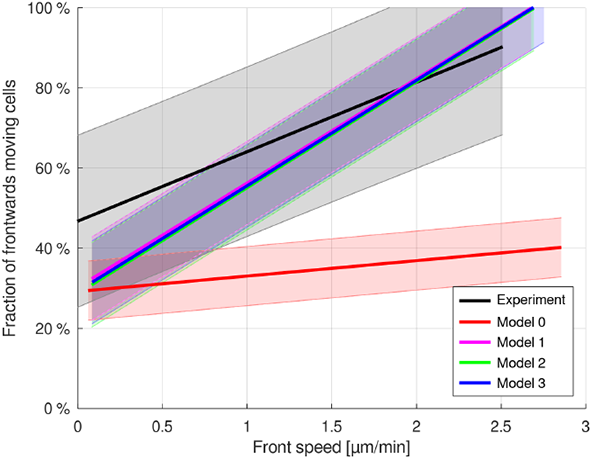
Uniformly distributed motility strength for polster cells 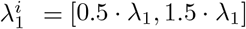 causes no significant changes in the simulations.

### 3.5 Robustness to time-varying motility parameters

As each polster cell can, when not guided, perform an individual run-and-tumble motion with gamma-distributed run duration and uniformly random orientation reset at tumble events, we also let the individual motility strength parameter 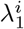 be drawn from a uniform distribution on the interval [0.5 *· λ*_1_, 1.5 *· λ*_1_] at gamma-distributed time points. Yet, no significant changes of cell movements could be found (cf. Fig. 8).

**Figure 8:**
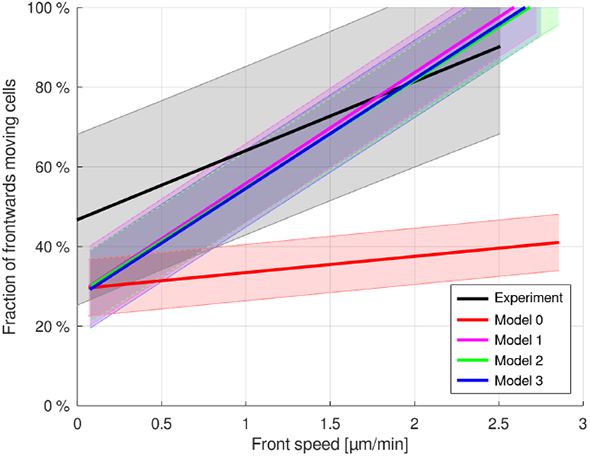
Randomly time-varying motility strength in the interval [0.5 *λ*_1_, 1.5 *λ*_1_] for all cells does not disturb the guidance-by-followers mechanism.

## 4 Discussion

The robustness of collective cell migration to variation is a classic question that falls within the broader framework of the robustness of developmental processes. Recently, we described a novel mechanism responsible for the collective guidance of polster cells and cohesion of the embryonic axis during zebrafish gastrulation [20]. The molecular details of guidance-by-followers are still not fully understood and, in particular, it is not clear how a cell integrates input from different neighbors. Here, we proposed different rule-based models with specific modalities of neighbor information integration by polster cells. We found that each of these models is able to recapitulate robust guidance of polster cells by axial mesoderm, suggesting that the guidance-by-followers process is robust to variation in cellular mechanisms. Although these results cannot indicate one solution that is closer to observations than another, we can at least support the idea that guidance-by-followers is a process sufficient to guide a group of cells regardless of how cells integrate information from multiple neighbors.

We then tested the robustness of these different models to variations in cell properties or in initial situation. First, we tested whether the ratio between the number of polster and axial cells affects collective guidance by running simulations with different ratios. We observed that, for each type of model, a large excess of polster cells leads to a disruption of collective guidance. In this case, the guidance cue and hence the collective behaviour in the polster is not sufficient to overcome the noise of individual cell behaviours. This result is particularly interesting as it suggests that there is a critical ratio threshold below which collective guidance is not possible, which is consistent with other studies reporting such a threshold for proper collective guidance [35, 36].

Cell shape and size is also known to be a critical factor in collective cell migration. In a confluent monolayer, regions of small cells and high density exhibit disordered movement, mainly due to steric interactions [37, 38]. Some studies also suggest that the intensity of juxtacrine signalling between cells is regulated by the cell-cell contact area [39, 40]. Thus, cell size may affect both contact-mediated guidance-by-follower signalling and cell mobility in a dense group. We tested the effect of cell size on collective guidance of the polster by introducing cell-to-cell variability. We observed that in the three models we proposed, the system is robust enough to cope with heterogeneity in cell size. Interestingly, this observation contradicts the only study that we are aware of on the effect of cell size during zebrafish gastrulation [41]. By manipulating ploidy, the authors created embryos with larger or smaller cells and observed mild gastrulation defects. In particular, they generated chimeric embryos with larger or smaller cells placed in WT embryos, and observed that the abnormally sized cells displayed similarly fast but less directed migration. However, this study does not focus on axial cells and it is not clear which cell population has been observed. Furthermore, it appeared that changing the size of a cell affected its cortical dynamics and thus its cell-autonomous migratory properties, whereas in our proposed model we assumed that apart from the variation in cell size, the rest of the cell properties were kept constant. Nevertheless, it would be interesting to test the prediction of this model by introducing larger and smaller cells in the axial mesoderm and measuring the effect on collective cell guidance.

Furthermore, we tested the robustness of the different models to variations in individual cell migration properties by varying how straight and fast polster cells tend to move during a run phase. Both heterogeneously distributed individual persistence and temporal variations both resulted in correct guidance of the cells, suggesting that this phenomenon is robust to differences in individual cell migration properties. This observation is particularly interesting as it suggests that, even in the case where individual polster cells are particularly poorly persistent, once they are set in motion by guidance-by-followers, they exhibit oriented collective behaviour. Such a phenomenon would ensure the robustness of axis development even in the case where the intrinsic guidance of polster cells is perturbed. Some molecular players are known to control cell migration persistence, such as Arpin or the CYFIP1/2 ratio in the WAVE complex [42, 43]. It would therefore be interesting to experimentally test whether local or global variations of individual cell persistence on polster cell guidance are buffered by guidance-by-followers.

In most simulations, although we observe a remarkable robustness of the model to parameter variation (except for an extreme cell type ratio), the slope of the oriented fraction as a function of axial mesoderm speed is always higher for simulated cells compared to experimental data. Since the individual migration behaviour of cells is well reproduced by the model, this observation suggests that there are some biological properties of guidance-by-followers that are not fully recapitulated by this basic model. However, it is interesting to note that such a simple set of rules is able to quasi quantitatively predict polster behaviour and is so robust to many variations. Also, in this study, we observed the robustness of the model to variations of a single parameter. It would be interesting to perturb several parameters together to further test the robustness of the model.

Finally, from a modeling perspective, this study highlights the role of individual-based approaches for understanding collective phenomena at the population scale that emerge from cell-cell interactions. Here, we have specifically assessed the robustness of our developed IBM to variations in modeling assumptions and to cell-to-cell variations. From a data management perspective, we ensure the reproducibility of this computational study and reusability and extensibility of the developed IBM by the strict separation of model definition from simulator code using the MorpheusML model description language [34] and a public model repository (https://morpheus.gitlab.io/model).

## Conflict of Interest Statement

The authors declare that the research was conducted in the absence of any commercial or financial relationships that could be construed as a potential conflict of interest.

## Author Contributions

RM and DJ developed the model, ran simulations, performed statistical analysis and prepared figures. JS developed, maintained and extended the model simulator. AB, ND and LB contributed to conception and design of the study. RM and LB wrote the first draft of the manuscript. All authors contributed to manuscript revision, read, and approved the submitted version.

## Funding

NBD acknowledges support by the French ANR, grant ANR-20-CE13-0016-LB acknowledges support by the German BMBF through EMUNE grant 031L0293D.

## Acknowledgments

We are grateful to Morpheus.lab members for fruitful discussions.

## Supplemental Data

Supplemental Movie 1 shows complete time courses of simulations from which snapshots were shown in Fig. 3. Supplemental Fig.S1 shows individual data points measured from simulations from which linear regression results were shown in Fig. 4.

## Data Availability Statement

The Morpheus model generated during this study was deposited in the public model repository under MorpheusModelID:M0008 (https://identifiers.org/morpheus/M0008). The model editor and simulator Morpheus is available open source at https://gitlab.com/morpheus.lab/morpheus under the BSD-3-Clause license. Pre-compiled packages are available for all major operating systems together with training materials and the model repository at the Morpheus homepage https://morpheus.gitlab.io.

## Notes

### Competing Interest Statement

The authors have declared no competing interest.

