## Supplementary figures and images for "Collective cell migration due to guidance-by-followers is robust to multiple stimuli"

### Supplemental Figure S1

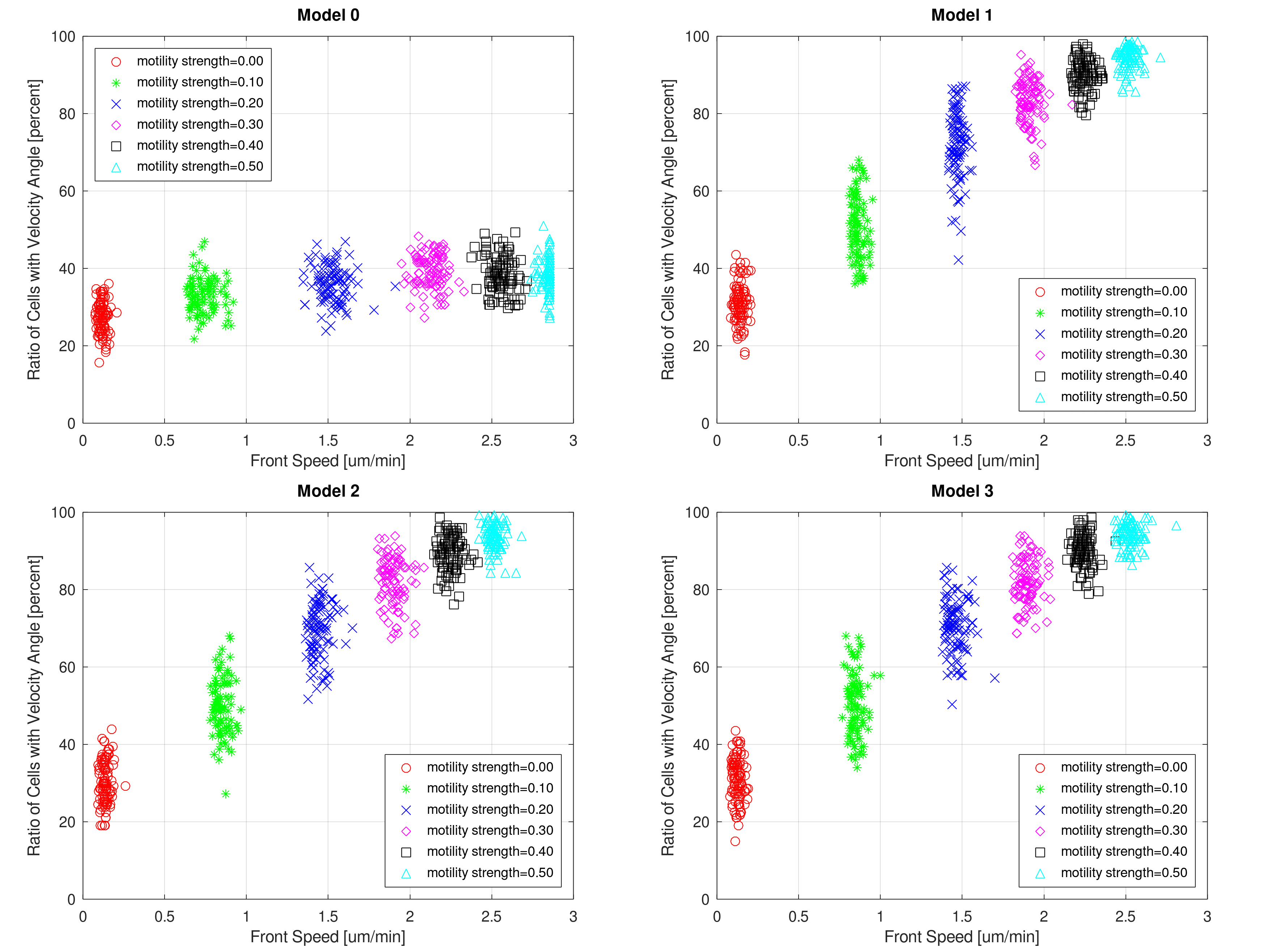
